# Quantitative estimates of deciduousness in woody species from a seasonally dry tropical forest are related to leaf functional traits and the timing of leaf flush

**DOI:** 10.1101/2021.03.03.433407

**Authors:** Souparna Chakrabarty, Sheetal Sharma, Shatarupa Ganguly, Asmi Jezeera, Neha Mohanbabu, Deepak Barua

## Abstract

Leaf phenology based classification of woody species into discrete evergreen and deciduous categories is widely used in ecology, but these categories hide important variation in leaf phenological behaviour. Few studies have examined the continuous nature of deciduousness and our understanding of variation in quantitative estimates of leaf shedding behaviour and the causes and consequences of this is limited. In this study we monitored leaf phenology in 75 woody species from a seasonally dry tropical forest to quantify three quantitative measures of deciduousness, namely: maximum canopy loss, duration of deciduousness, and average canopy loss. Based on proposed drought tolerance and drought avoidance strategies of evergreen and deciduous species, respectively, we tested whether the quantitative measures of deciduousness were related to leaf functional traits. Additionally, to understand the functional consequences of variation in deciduousness we examined relationships with the timing of leaf flushing and senescing. We found wide and continuous variation in quantitative measures of deciduousness in these coexisting species. Variation in deciduousness was related to leaf function traits, and the timing of leaf flushing. Along a continuous axis ranging from evergreen to deciduous species, increasing deciduousness was associated with more acquisitive leaf functional traits, with lower leaf mass per area and leaf dry matter content, and greater leaf nitrogen content. These results indicate that the continuous nature of deciduousness is an important component of resource acquisition strategies in woody species from seasonally dry forests.

## Introduction

Leaf phenology, the timing of leaf flushing, maturation and senescence, allows plants to match leaf growth, development and function to favourable environmental conditions. Additionally, leaf phenology plays an important role in interspecies, and trophic interactions (Van Schaik *et al*. 1993), and in determining seasonal patterns of ecosystem productivity and function (Wu *et al*. 2016, Xia *et al*. 2015). The use of leaf phenology based classification to group woody plants into discrete evergreen and deciduous categories has a long history, and has been broadly applied in ecology (Axelrod 1966). This categorization has been useful in providing insight into shared physiological and structural adaptations, and common responses to environmental change and disturbance (Givnish 2002, Kikuzawa & Lechowicz 2011). However, despite this long tradition of use and broad utility, these discrete categories hide potentially important variation in leaf phenological behaviour, and few studies have examined the continuous nature of the extent of deciduousness (but see Kushwaha *et al*. 2010, Williams *et al*. 2008). In this study we monitored leaf phenology in woody species from a seasonally dry tropical forest to quantify deciduousness and examined relationships of quantitative estimates of deciduousness with leaf functional traits and the timing of leaf phenological activity.

Woody species in seasonally dry tropical forests (STDF) display a wide diversity of leafing behaviours (Borchert *et al*. 2002; Eamus 1999; Kushwaha *et al*. 2010; Rossatto *et al*. 2012; Singh & Kushwaha 2005; Williams *et al*. 1997; Williams *et al*. 2008). At the two extremes, evergreen species maintain most of their canopy through the year, while in contrast, deciduous species shed leaves and remain leafless for some duration in the dry season (Eamus 1999). There are a range of species that display an intermediate behaviour and lose some fraction of their canopy during the dry season, and these are commonly classified as semi-evergreen, semi-deciduous, or brevideciduous species (Borchert *et al*. 2002; Eamus 1999; Singh & Kushwaha 2005; Williams *et al*. 2008). While these finer categories try to capture variation in the extent of leaf loss, the terminology has been inconsistently used with different terms used to describe similar behaviours, or different and somewhat arbitrary thresholds of canopy loss used to define these subcategories (Singh & Kushwaha 2005). Additionally, for species that are deciduous, the duration that they remain leafless can vary substantially from less than a few weeks to several months (Borchert *et al*. 2002; Kushwaha & Singh 2005; Williams *et al*. 2008). Moreover, the classification into discrete evergreen and deciduous categories is complicated by intra-specific variation in leaf shedding behaviour as the complete shedding of leaves in some species can vary across individuals in a population as a function of spatial heterogeneity in water availability, and across years as a consequence of varying water availability. Such variation in the magnitude and duration of deciduousness is likely related to morphological and physiological traits, and is expected to have important consequences for resource acquisition.

Despite the wide variation documented, few studies have attempted to quantify this potentially important trait. Williams *et al*. (2008) used two metrics, maximum canopy loss (MCL) and duration of deciduousness (DD) to examine variation in deciduousness in woody species from a seasonally dry forest in Thailand. MCL, a measure of the magnitude of deciduousness, ranged from species that maintained most of their canopy through the year to those that lost all their leaves. DD ranged from zero for the evergreen species to five months for the very deciduous species. While this study did not go much beyond describing the two quantitative measures used to estimate deciduousness, one of the noteworthy results was the near continuous variation in the nature of deciduousness reported in these coexisting species. The duration that species remain leafless has been quantified more often in other studies (Borchert *et al*. 2002; De Bie *et al*. 1998; Eliot *et al*. 2006; Kushwaha & Singh 2005; Olivares & Medina 1992; Rivera *et al*. 2002). These studies show wide variation in the duration of deciduousness from very short periods of less than a week in leaf exchanging species to as high as seven months in very deciduous species. The utility of these two measures, the magnitude and duration of deciduousness may differ for evergreen and deciduous species. While the magnitude of deciduousness is useful to distinguish between evergreen species that lose different fractions of their canopy, it cannot differentiate between deciduous species that lose their entire canopy during the dry season. Conversely, the duration of deciduousness is useful to distinguish between deciduous species that remain leafless for different durations, but not for evergreen species that are never completely leafless. Another measure that has not received much attention (but see Sastry & Barua 2017) is an estimate of the annual canopy maintained (or lost) through the year. This measure has the advantage of integrating information from both the extent and duration of deciduousness and can be used to quantify the leafing behaviour of both evergreen and deciduous species (Fig. 1).

**Figure 1:**
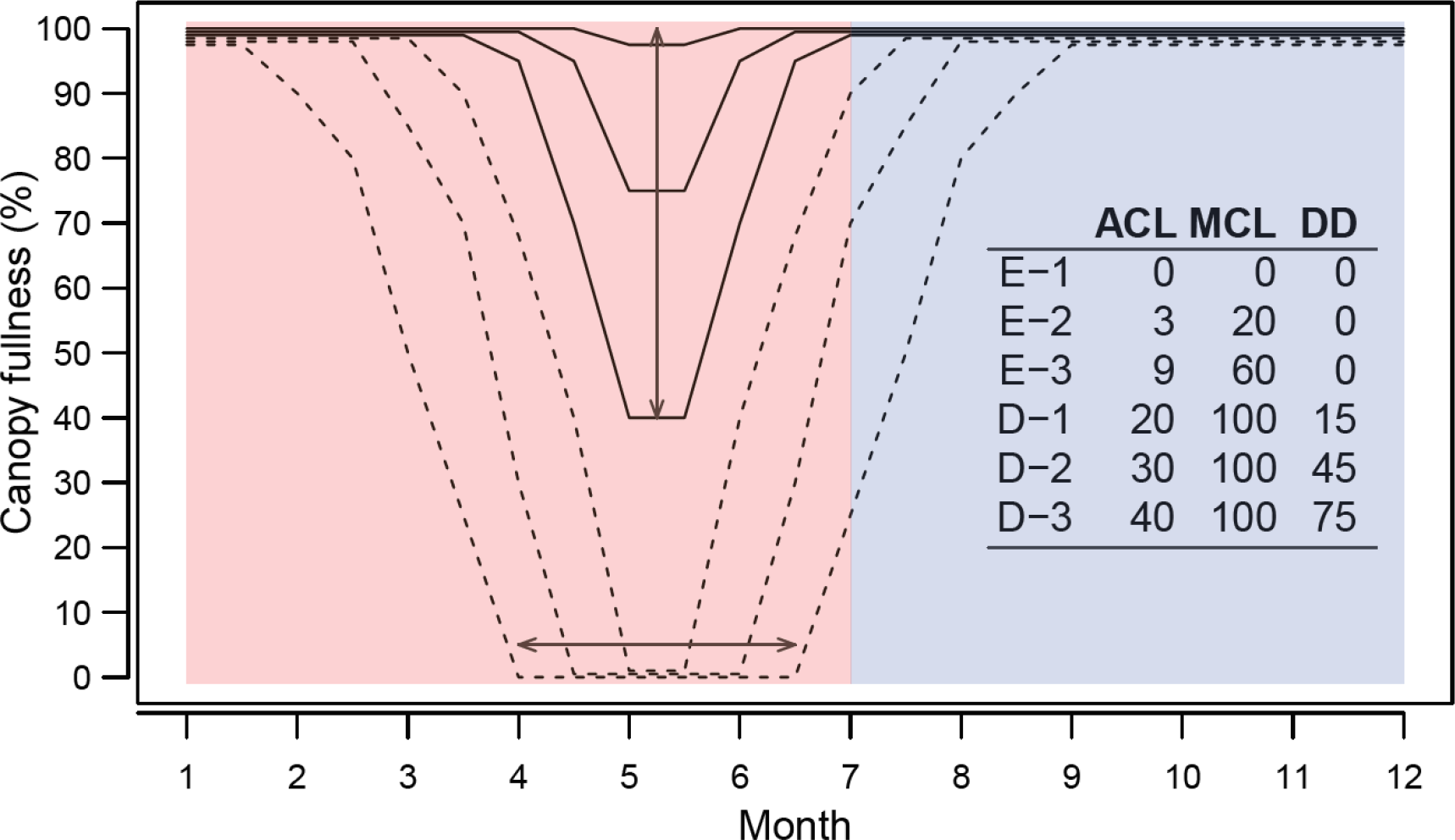
Conceptual representation of canopy fullness through the season and variation in leaf shedding behaviour in three hypothetical evergreen (solid lines, E-1 to E-3) and deciduous species (dashed line, D-1 to D-3) species. The vertical line with double arrows depicts the maximum loss of canopy (MCL) for one of the evergreen species, and the horizontal line with double arrows depicts the duration of deciduousness (DD) for one of the deciduous species. The table in the inset provides estimates for average canopy loss through the year (ACL), maximum canopy loss (MCL), and duration of deciduousness in days (DD) for these six hypothetical species. The dry and wet season are indicated by the red and blue colours respectively.

In seasonally dry tropical forests (STDF), water is typically abundant during the wet season, but water limitations during the dry season impose severe constraints on plant function, growth and survival (Eamus 1999, Murphy 1986). Plants in these forests must endure the unfavourable conditions during the dry season, but also maximize growth and productivity during periods when water is more abundant. Two alternate strategies of drought tolerance and drought avoidance have been proposed to cope with the seasonal availability of water (Givnish 2002; Markesteijn & Poorter 2009; Mendez-Alonzo *et al*. 2012). Evergreen species adopt drought tolerant strategies and retain most of their canopy through the year. This allows photosynthetic activity through most of the year, but to be able to sustain this, these species must possess adaptations to withstand low soil water potentials, minimize water loss, and improve water use efficiency (Sterck *et al*. 2011; Wright, Reich & Westoby 2001; Wright, Westoby & Reich 2002). In contrast, drought avoiders shed leaves early in the dry season, likely to prevent water loss (Borchert *et al*. 2002, Tyree *et al*. 1993; Mendez-Alonzo *et al*. 2012). These deciduous woody species do not have to invest in traits related to enduring low soil water or minimizing water loss, and this allows them to adopt exploitative or acquisitive resource use strategies that maximize growth and photosynthesis when water is available (Mendez-Alonzo *et al*. 2012; Mendez-Alonzo *et al*. 2013; Sterck *et al*. 2011).

The distinct acquisitive and conservative strategies in woody species in STDF imply that it may not be possible to maintain leaves through the year, and concurrently maximize growth and productivity during favourable conditions. Indeed, leaf and stem functional traits in evergreen and deciduous species suggest that such a tradeoff may be mediated via leaf and stem functional traits. Evergreen species typically have higher leaf mass per area (LMA) and leaf dry matter content (LDMC), but lower leaf nitrogen content (LNC) (Poorter *et al*. 2009; Wright *et al*. 2005; Mendez-Alonzo *et al*. 2012). In contrast, deciduous species have lower LMA and LDMC, and higher LNC. Additionally, evergreen species from STDF have been reported to have higher wood density with lower hydraulic efficiency, but higher resistance to embolism (Mendez-Alonzo *et al*. 2012). This is in contrast to lower density wood with higher hydraulic efficiency, but greater sensitivity to embolism in deciduous species (Mendez-Alonzo *et al*. 2012). Thus, broad-leaved evergreen and drought deciduous species in STDF may represent two extremes along the leaf and whole plant economic spectrums indicative of conservative and acquisitive strategies, respectively. However, we do not know if variation in the magnitude and duration of leaf loss in species from these discrete leaf habit categories is related to plant functional traits.

While peak leaf flushing in the dry season is widely reported in seasonally dry forests, there is substantial variation in the timing of flushing among species (Singh & Kushwaha, 2005; Elliott *et al*., 2006; Williams *et al*., 2008; Borchert, 1994; Eamus, 1999). Typically, evergreen species that can withstand the low soil moisture and high vapor pressure deficit during the dry period flush leaves earlier in the dry season. This allows evergreen species to utilize the higher light availability in the drier parts of the year to maximize photosynthesis, growth and reproduction (van Schaik, Terborgh & Wright 1993; Eamus 1999). In contrast, deciduous species typically flush closer to the onset of the rains, and shed leaves early during the dry season in response to the decreasing soil water availability and increasing VPD (Elliott, Baker & Borchert 2006, Williams-Linera 1997; Kushwaha & Singh 2005). If quantitative estimates of deciduousness are functionally relevant, we would expect to see a relationship with the timing of leaf flushing with more evergreen species flushing earlier and the more deciduous species flushing closer to the onset of the rains. Similarly, we would expect to see a relationship with leaf senescence, with the more deciduous species shedding leaves earlier in the dry season.

We monitored leaf phenology in 75 species from a seasonally dry tropical forest to quantify the seasonal loss of leaves. For these species, we quantified three measures of deciduousness: 1) Maximum canopy loss (MCL); 2) Duration of deciduousness (DD); and, 3) Average annual canopy loss (ACL). We asked if these continuous measures of deciduousness were related to leaf functional traits, namely leaf mass per area (LMA), leaf dry matter content (LDMC), leaf area (LA), leaf carbon content (LCC) and leaf nitrogen content (LNC). Based on known differences in leaf functional traits between evergreen and deciduous species, and the relationship of leaf functional traits with tolerance to abiotic conditions and leaf longevity, we expected that measures of deciduousness would be negatively related to LMA, LDMC, LA and LCC; and positively related to LNC. To understand the functional implications of the measures of deciduousness we tested if they were related to the timing of leaf flushing and senescence. We predicted that species with higher deciduousness would flush later in the year closer to the rains, while the more evergreen species with lower estimates of deciduousness would flush earlier to match peak photosynthetic performance with maximum light availability. We also expected species that are more deciduous to be more sensitive to seasonal drought and shed leaves earlier in the dry season, than the less deciduous (more evergreen) species.

### Materials and methods

#### Study site

This study was conducted in the Northern Western Ghats, around Nigdale, Maharashtra, India 19°21’ -19°11’N, 73°31’ -73°37’E, 900 m asl. The region lies in the Northern extremes of the Western Ghats range. The landscape consists of valleys carved out by the river Bhima and its tributaries. The top of the valleys consists of flattened ridges. Soil depth varies from shallow (< 10cm) at the top of these ridges to deep (>100cm) in the mid slopes and valleys (Ghadage *et al*. 2013). The vegetation is heterogeneous and varies from open savanna type woodlands in the ridge tops to closed-canopy forests in the valleys. The vegetation differs markedly based on topographic and edaphic conditions. The open forests on the ridge tops have a very low soil depth, low water availability, high light availability, and 30-70 percent tree cover. The woody vegetation in open habitats has a high percentage of deciduous species consisting of stunted trees (3-5 m), lianas and shrubs. At the other extreme the closed valley forests are characterized by greater soil depth, higher soil water availability, and low light availability in the understory. These habitats are dominated by tall statured evergreen trees (15-25 m), lianas and under canopy shrubs. Habitats between the open crests and the valleys consist of smaller fragmented patches of forests along the slopes where soil depth varies between 10-80 cm. This transition zone contains a mixture of evergreen and deciduous species with some overlap with both open and closed habitats. Rainfall is highly seasonal and most of the annual average rainfall of 2266 mm falls between June and September. Rainfall between December and May is minimal and averages less than 30 mm per month. Minimum temperatures in January average 11 °C while maximum temperatures in April and May average 35 °C. The climate data were obtained from the CRU 2.0 dataset (New *et al*. 2002).

#### Phenology monitoring

Vegetative phenology was monitored monthly for a period of 48 months from January 2014 to December 2017. Phenology censuses were conducted by direct visual observation with the aid of binoculars around the 15^th^ of every month and typically completed in a period of 3 to 4 days. We monitored a total of 1303 individuals of 75 species. We targeted a minimum of 10 individuals per species, but availability for some rare species was limited and the number of individuals monitored ranged from 7 to 8 for three species. The number of individuals monitored ranged 10 to 14 for 21 species that were not very common, and greater than or equal to 15 the remaining 51 species. Phenology observations were initiated a year prior to the final study duration and this time was used to calibrate and standardize estimates for each species. All phenology observations were conducted by the same observer to avoid observer bias. The total canopy fullness of individuals was scored in a semi-quantitative manner on a scale from 0 to 100 %, where a score of 0 represents the complete absence of leaves, and a score of 100 represents a full canopy. Additionally, the canopy for every individual was scored for the proportion of leaves that were flushing, mature and senescing based on size, colour and texture of leaves.

Three quantitative measures were used to characterize the nature of deciduousness. These include: 1) Maximum canopy loss (MCL) - the maximum loss of canopy observed in a year; 2) Duration of deciduousness (DD) - the duration in days that an individual was completely leafless in a year; and, 3) Average canopy loss (ACL) - an integrated measure of the average canopy loss observed through the year. MCL, DD and ACL were calculated for every individual per year, and averaged across all individuals and years to get a representative value for each species. At the level of species, we defined a species as potentially deciduous when at least 25 % of the monitored individuals were completely leafless for any of the years. Principal component analysis was performed for the three measure of deciduousness, MCL, DD and ACL using the R package ’factoextra’ (Kassambara & Mundt 2020).

For determining the timing of leaf flushing and senescing we first estimated the activity period for every species. This was defined as the earliest and latest month across all years of observation when at least 20% of individuals had started and stopped leaf flushing and senescing activity, respectively. As activity periods for species did not follow the calendar year, *i*.*e*., did not necessarily start in January and end in December, we have complete activity periods for only three of the four years in the study duration. This limited the number of years for which we could estimate measures of deciduousness, and the timing of flushing and senescing to three years. Within the activity period for each species, circular statistics was used to calculate the vector length for flushing and senescing activity for every individual by weighing the date of census expressed as an angle representative of the day of the year by the respective flushing and senescing activity score on those dates (Zar 1999). Rayleigh’s test for uniformity was conducted on vector lengths to determine if the distribution of flushing and senescing activity was uniform. If different from a uniform distribution, a mean angle representing peak flushing and senescing activity was calculated using circular statistics (Zar 1999). These were averaged across all individuals and years to determine a mean flushing and senescing date for each species. Angular-linear correlation analyses were used to examine relationships between quantitative measures of deciduousness and species mean flushing and senescing dates using the R package ’Directional’ (Tsagris *et al*. 2020).

#### Trait quantification

We followed protocols recommended by Perez-Harguindeguy *et al*. (2013) for the collection and processing of samples to quantify leaf functional traits. Three to six fully expanded and mature leaves were collected from the sun-exposed upper canopy of three to six individuals of every species. For understory species where sun-exposed leaves were not available, we collected leaves from the upper canopy. A combination of a telescopic leaf pruner with an 8 m reach, and professional tree climbers were used to access the upper canopy. Leaves were placed in sealed plastic bags which contained moistened paper towels to keep the air in the bags water saturated and prevent desiccation of the leaves. Samples were transported back to the laboratory on the same day, and water saturated overnight by immersing the petiole in a container filled with water. Water saturated leaves were weighed to determine saturated fresh weight, scanned with a desktop scanner for quantifying leaf area (LA), and then oven-dried at 60 °C for 72 hours to determine dry weight. Leaf mass per unit area (LMA) was quantified as the ratio of dry weight to area, and leaf dry matter content (LDMC) as the ratio of dry weight to saturated fresh weight. Leaves for quantification of LMA, LA and LDMC were collected from all 75 study species between 2013 and 2015. A separate collection was initiated between 2015 to 2016 to quantify leaf carbon and nitrogen content. For a subset of 55 of the dominant species in this community, five leaves from at least five individuals for each species were collected as described before, oven dried at 60 °C for 72 hours and then ground to a fine powder to quantify leaf carbon concentration (LCC) and leaf nitrogen concentration (LNC) using a LECO CHN Elemental Analyzer (Truspec Micro CHNS, LECO, USA). Leaf carbon and leaf nitrogen were quantified as a percent of unit dry mass. Pearson’s correlations were used to examine relationships between quantitative measures of deciduousness and leaf functional traits.

## Results

In addition to the wide variation observed, one of the striking results was the near continuous nature of variation in the measures of deciduousness (Fig. 2). The maximum loss of canopy (MCL) ranged from near zero to 100 %. For the evergreen species MCL ranged from zero to greater than 70 %, and for deciduous species from around 40 to 100 %. Estimates of maximum canopy loss (MCL) were not 100 % for deciduous species due to intra-population variation in deciduousness resulting from intra-individual and inter-annual variation in leaf loss behaviour. The duration of deciduousness ranged from near zero for most evergreen species to greater than 100 days in the very deciduous species. Average canopy loss (ACL) ranged from near zero for some of the evergreen species that maintained most of their canopy through the year to near 50 % for the most deciduous species. For the evergreen species ACL ranged from zero to greater than 25 %, and for deciduous species from as low as 6 % to greater than 50 %.

**Figure 2:**
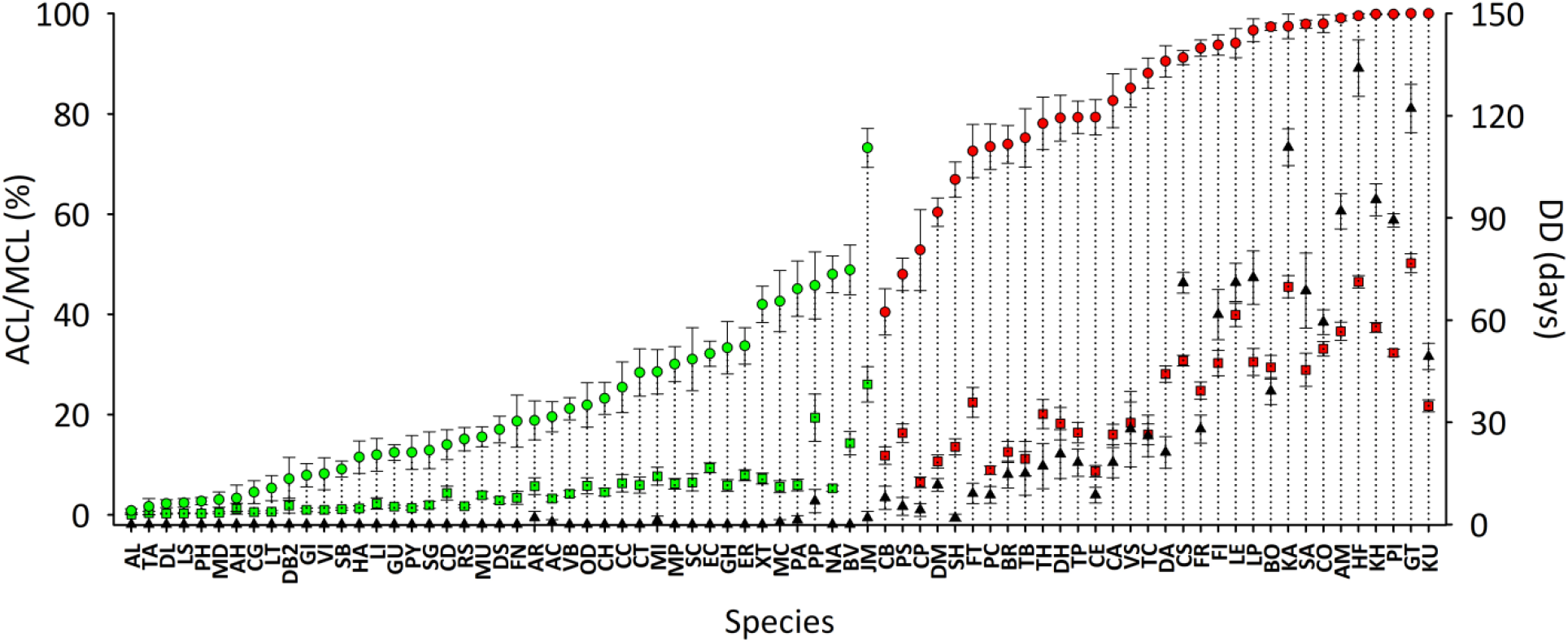
Average canopy loss (ACL, squares), maximum canopy loss (MCL, circles) and duration of deciduousness (DD, triangles) in the study species. For ACL and MCL evergreen species are depicted in green and deciduous species in red. ACL and MCL are presented as a percentage and DD in days. Error bars represent standard errors. Species names corresponding to the abbreviations used are available in Table S1.

The continuous measures of deciduousness were strongly correlated to each other (Fig. 3). However, measures of maximum canopy (MCL) loss failed to separate the deciduous species while deciduous duration (DD) was not useful for differentiating between evergreen species (Fig. 3c).

**Figure 3:**
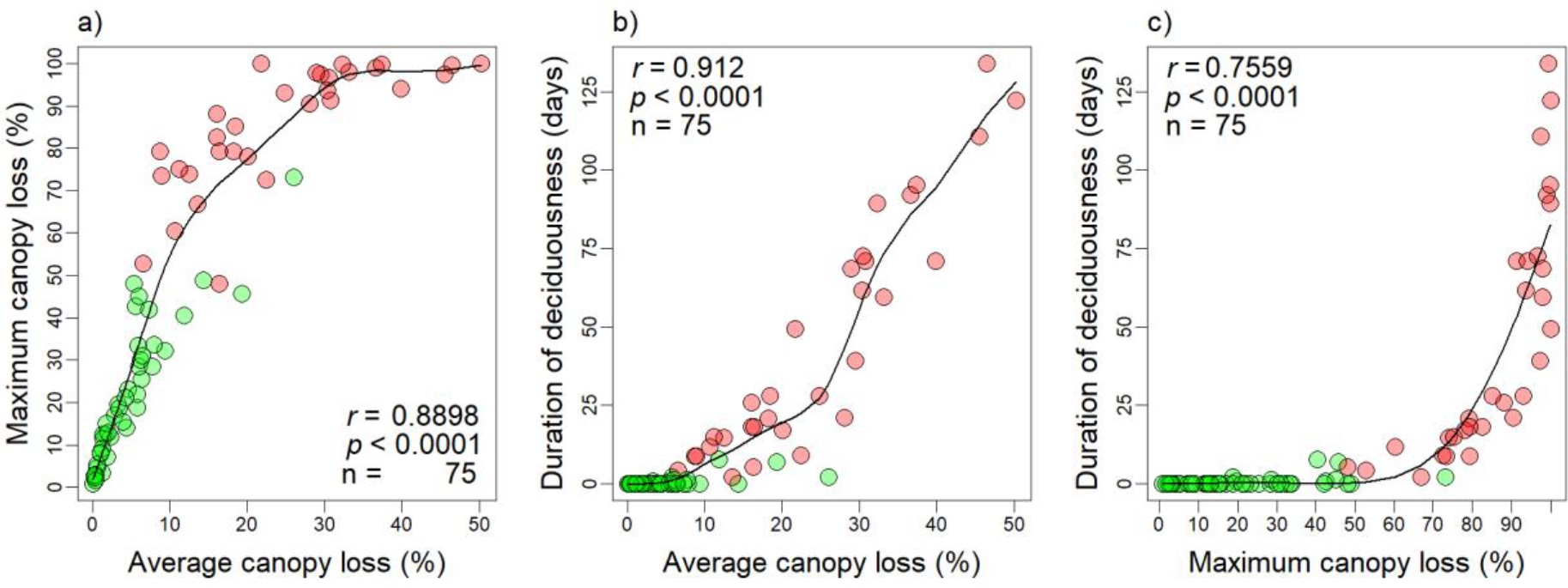
Relationships between the three measures of deciduousness: Average canopy loss (ACL), maximum canopy loss (MCL) and duration of deciduousness (DD) for the study species. Evergreen species are depicted in green and deciduous species in red. The black solid line is a spline fit presented only as a visual aid. Pearson’s correlation coefficients, tests for significance and sample size are presented in each panel.

Congruent with the strong relationships between these measures of deciduousness, principal component analysis revealed that most of the variation in these three measures was explained by the first principal component axis (Fig. 4). Average canopy loss (ACL) showed the highest loading on the first principal component axis. Following these results, we concentrated on ACL to examine relationships between deciduousness and leaf traits and timing of leaf flushing and senescence.

**Figure 4:**
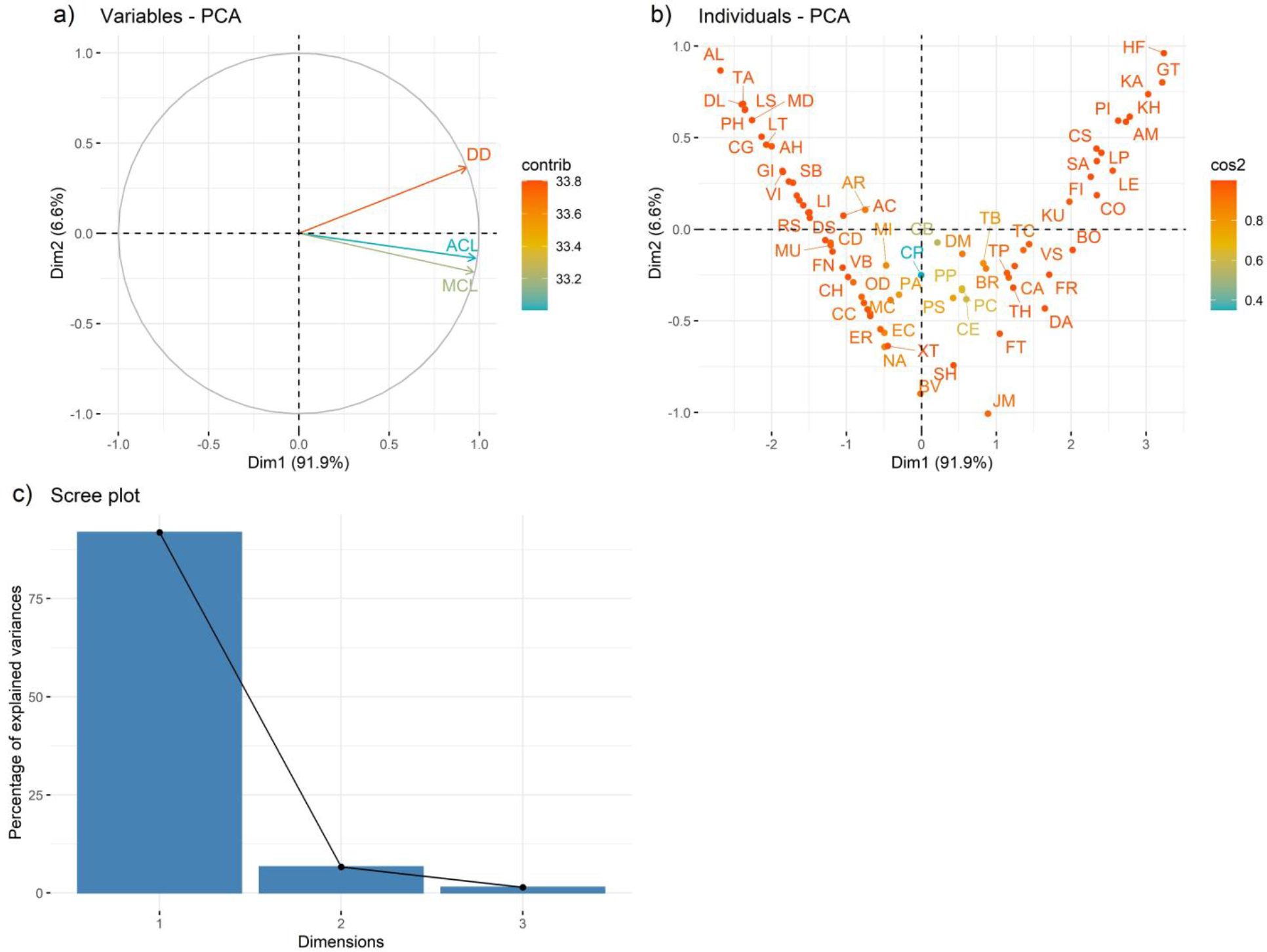
Principal component analysis of the three measures of deciduousness: maximum canopy loss (MCL), duration of deciduousness (DD), and average canopy loss (ACL). Biplot for principal components 1 and 2 depicting: a) the three measures of deciduousness; and b) showing the individual species. Panel c) shows the Scree plot with the percentage of variance explained by principal components 1, 2 and 3. Species names corresponding to the abbreviations used are available in Table S1.

Measures of deciduousness were negatively related to leaf mass per area, leaf dry matter content, leaf C:N ratio, and positively related to leaf nitrogen content (Fig. 5, Fig. S1). We did not observe any significant relationships between measures of deciduousness and either leaf area or leaf carbon content (Fig. S1).

**Figure 5:**
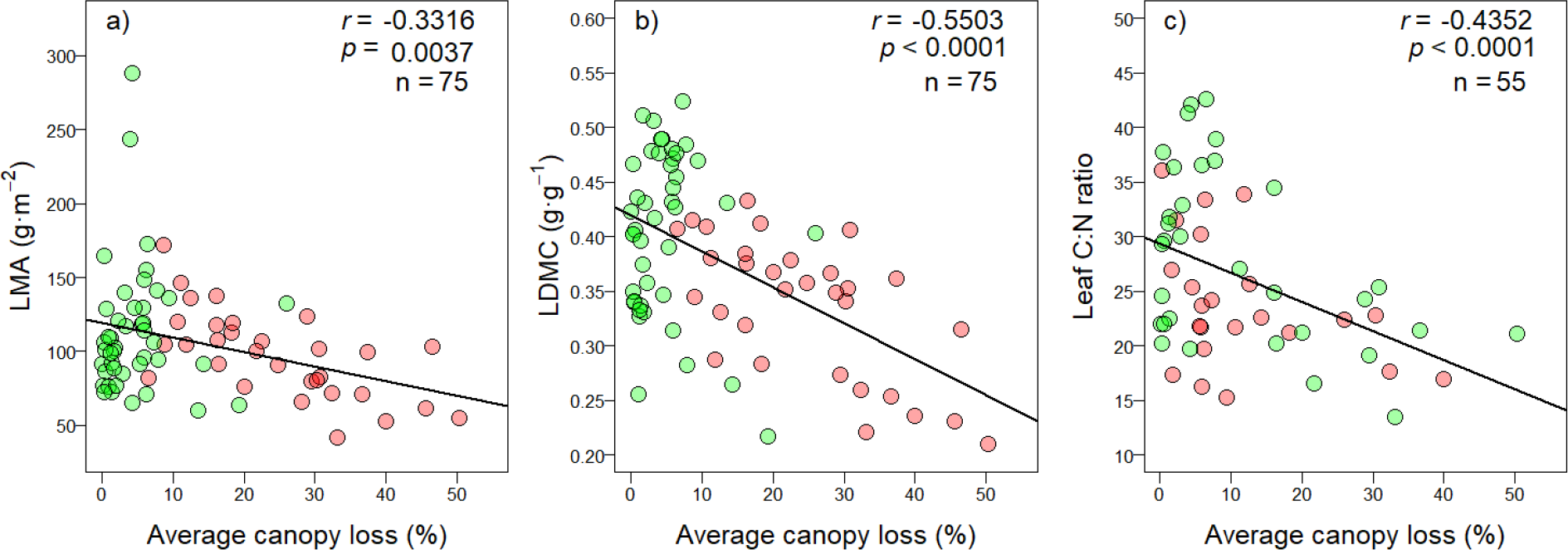
Relationship between average canopy loss (ACL) as a quantitative measure of deciduousness and leaf functional traits. Deciduous species are depicted in red and evergreen species in green. Pearson’s correlation coefficients, tests for significance and sample size are presented in each panel, and regression lines are shown as a visual aid.

Mean flushing timing of these coexisting species was spread out across the year (Fig. 6a). Most evergreen species flushed their leaves in the early dry season. In contrast, deciduous species flushed leaves closer to the onset of the rains. Additionally, the most deciduous species with highest average canopy loss flushed leaves after the onset of the rains. Mean species flushing time was significantly related to average canopy loss (r^2^_angular-linear_ = 0.7086, p < 0.001) and other measures of deciduousness (Table S2) confirming that more deciduous species with higher ACL scores flushed closer to or after the onset of the rains. Mean senescence time for species were more clustered seasonally with most species shedding leaves in the early part of the dry season (Fig. 6b). In contrast to the timing of leaf flushing, there were no obvious differences in the timing of leaf senescence between evergreen and deciduous species. Congruent with this we did not find a significant relationship between mean species senescence time and average canopy loss (r^2^_angular-linear_ = 0.09, p > 0.05). Mean senescence timing was correlated to MCL and DD but these relationships were weaker than what we observed with ACL (Table S2).

**Figure 6:**
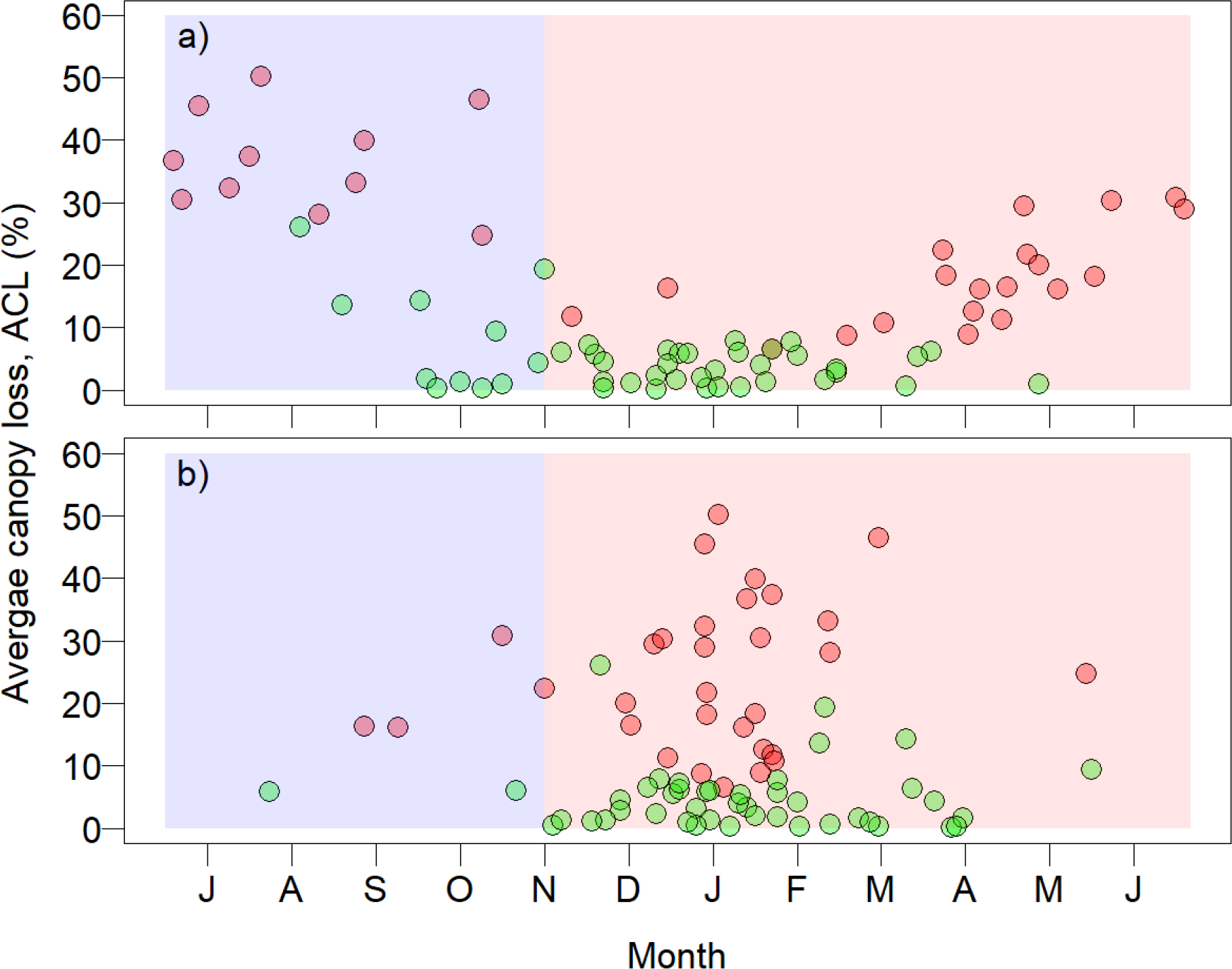
Relationship between average canopy loss (ACL), a quantitative measure of deciduousness and the timing of: a) leaf flushing, and b) leaf senescing. Deciduous species are depicted in red and evergreen species in green. The seasonal timeframe chosen here run from the beginning of the rainy season (depicted in blue) in July to the end of the dry season (depicted in red) in June.

## Discussion

Interspecific variation in leaf phenology is an important axis of functional diversity in plants. Phenology-based classification into discrete evergreen and deciduous categories has been broadly applied in ecology and has provided fundamental insights into shared physiological and structural adaptations, and common responses to environmental change and disturbance. However, few studies have examined the continuous nature of the extent of deciduousness. Results from this study show that these discrete categories hide important variation in continuous estimates of leaf loss behaviour. This variation in continuous measures of deciduousness in coexisting species from a seasonally dry tropical forest was related to leaf functional traits indicating that quantitative estimates of the intensity and duration of leaflessness constitutes an important component of resource acquisition strategies. Finally, measures of deciduousness were functionally important and related to the timing of leaf flushing.

The range of maximum canopy loss (MCL) and duration of deciduousness (DD) observed in these 75 species was similar to what has been reported in the few studies that have examined this (Borchert *et al*. 2002; De Bie *et al*. 1998; Eliot *et al*. 2006; Kushwaha & Singh 2005; Olivares and Medina 1992; Rivera *et al*. 2002; Williams *et al*. 2008). In addition to these two metrics, we used average canopy loss (ACL) as novel measure for estimating deciduousness. ACL integrates information from both DD and MCL, overcomes shortcomings of MCL and DD, and can be used effectively for evergreen and deciduous species. As seen in our results, MCL was effective in differentiating between evergreen species but not deciduous species that lose their entire canopy. In contrast, DD was effective in differentiating between deciduous species but not evergreen species that are never completely leafless. Not surprisingly, these measures of deciduousness were strongly correlated to each other, and principal component analyses revealed that most of the variation in these three measures was explained by the first principal component axis with ACL showing the highest loading on the first principal component axis. Based in these results we suggest that ACL, an intuitive continuous measure of deciduousness that can be estimated from ground-based phenology studies is a useful quantitative index for deciduousness.

Evergreen and deciduous species in STDF are thought to represent distinct alternate strategies of drought tolerance and drought avoidance, respectively (Givnish 2002; Markesteijn & Poorter 2009; Mendez-Alonzo *et al*. 2012).The drought tolerant strategies of evergreen species are associated with conservative leaf and stem traits related to adaptations to withstand low soil water potentials, minimizing water loss, and improving water use efficiency (Sterck *et al*. 2011; Wright, Reich & Westoby 2001; Wright, Westoby & Reich 2002). In contrast, the avoidance strategies of deciduous species are associated with traits that allow these species to maximize resource acquisition when water is abundant (Mendez-Alonzo *et al*. 2012; Sterck *et al*. 2011). We observed significant relationships between our continuous measures of deciduousness and leaf functional traits. Along a continuous axis ranging from evergreen to deciduous species, increasing deciduousness was associated with more acquisitive leaf functional traits, with lower leaf mass per area and leaf dry matter content, and greater leaf nitrogen content. These results indicate that leaf phenology, particularly the continuous nature of deciduousness is intricately linked to resource acquisition strategies in woody species from seasonally dry forests.

The quantitative estimates of deciduousness were functionally important as indicated by the significant relationships with the timing of leaf flushing. Most evergreen species flushed their leaves early in the dry season, while most deciduous species flushed leaves later in the dry season closer to the onset of the rains. Interestingly, the most deciduous species in this community with the highest ACL scores flushed leaves after the onset of the rains during the rainy season. Thus, on the continuous scale of deciduousness, less deciduous species may be able to flush earlier in the dry season because conservative leaf functional traits allow them to maintain function despite the low soil moisture availability and high vapour pressure deficits experienced through the dry season (Poorter *et al*. 2009; Wright *et al*. 2005; Mendez-Alonzo *et al*. 2012). Species that are more deciduous have acquisitive leaf functional traits associated with greater sensitivity to low water availability (Mendez-Alonzo *et al*. 2012; Sterck *et al*. 2011), and flush leaves closer to the onset of the rains. The species with the highest ACL scores in these forests were those that flushed after the onset of the rains. However, relationships between measures of deciduousness and the timing of leaf senescence were weaker and observed for MCL and DD, but not ACL. Overall, leaf senescence was aggregated in time indicating that the timing of leaf senescence may be constrained due to strong selection by abiotic conditions. The limited variation in the timing of senescence may explain the weaker relationship with quantitative measures of deciduousness.

Drought deciduousness is thought to be a mechanism that helps in the conservation of water and protect plants from reaching critically low water potential that may result in catastrophic hydraulic failure (Tyree *et al*., 1993). However, there is little empirical evidence to support this widely held assumption, and it has been suggested that the role of drought deciduousness as a water conservation strategy may not be universal (Wolfe *et al*. 2016). The observed relationship between continuous measures of deciduousness and leaf functional traits suggests an alternate explanation for drought deciduousness in woody species. Increased deciduousness is related to an overall reduction in the potential carbon gain over the growing season. Increased resource acquisition from exploitative strategies in deciduous species can make up for this loss in carbon gain. Traits associated with acquisitive strategies are inherently negatively related to leaf life span and sensitivity to harsh environmental conditions (Mendez-Alonzo *et al*. 2012; Sterck *et al*. 2011; Wright *et al*. 1994). Thus, shifts towards more acquisitive strategies that allow greater carbon gain over shorter durations when water is more abundant are likely intrinsically linked to shorter leaf life spans and lower tolerance to drought. This suggests that drought deciduousness may be a consequence of acquisitive strategies rather than a mechanism for water conservation.

Overall, these results highlight the importance of continuous variation in deciduousness in woody species. In addition to maximum canopy loss and duration of deciduousness, average canopy loss is a novel, intuitive, and readily measurable index can be to effectively used to estimate the continuous nature of deciduousness in woody species. These measures of deciduousness are related to leaf functional traits, are functionally important in being related to the timing of leaf flushing, and useful in providing a mechanistic understanding of species responses to seasonal environments.

## Acknowledgements

Authors acknowledge assistance in the field from Kalu, Ganpat and Thakkubai; Mahesh Sankaran for facilitating the processing of the leaf carbon and nutrient analysis: IISER Pune for intramural funding and support; and SERB for funding.

## Author contributions

DB designed the study; SC carried out the phenology monitoring; SS, SG, AJ and NM quantified leaf functional traits; SC and DB analyzed the data and wrote the manuscript.

