## Supplementary Material for "Quantitative estimates of deciduousness in woody species from a seasonally dry tropical forest are related to leaf functional traits and the timing of leaf flush"

**Contents:**

**Table S1:** The study species with taxonomic affiliation (Family), species code (Code), life form (Life), leaf habit (Leaf), and number of individuals monitored (n).

**Table S2:** Relationship between measures of deciduousness and species mean flushing and senescing dates.

**Figure S1:** Correlation matrix for quantitative measures of deciduousness and leaf functional traits.

**Table S1:** The study species with taxonomic affiliation (Family), species code (Code), life form (Life), leaf habit (Leaf), and number of individuals monitored (n). Abbreviations: Life form: T = Tree, L = Liana, and S = Shrub; Leaf habit: E = Evergreen, and D = Deciduous.

| | Species | Family | Code | Life | Leaf | n |
|---|---|---|---|---|---|---|
| 1. | Acacia concinna (Willd.) DC. | Leguminosae | SH | T | D | 15 |
| 2. | Actinodaphne gullavara (Buch.-Ham. ex Nees) M.R.Almeida | Lauraceae | AC | T | E | 27 |
| 3. | Aglaia lawii (Wight) C.J.Saldanha | Meliaceae | AL | T | E | 24 |
| 4. | Allophylus cobbe (L.) Raeusch. | Sapindaceae | TP | S | D | 33 |
| 5. | Ancistrocladus heyneanus Wall. ex J.Graham | Ancistrocladaceae | HA | L | E | 11 |
| 6. | Artocarpus heterophyllus Lam. | Moraceae | AH | T | E | 7 |
| 7. | Atalantia racemosa Wight ex Hook. | Rutaceae | AR | T | E | 15 |
| 8. | Bridelia retusa (L.) A.Juss. | Phyllanthaceae | BR | T | D | 30 |
| 9. | Caesalpinia cucullata Roxb. | Leguminosae | MC | L | E | 14 |
| 10. | Callicarpa tomentosa (L.) L. | Lamiaceae | CT | T | E | 14 |
| 11. | Carallia brachiata (Lour.) Merr. | Rhizophoraceae | CB | T | E | 14 |
| 12. | Careya arborea Roxb. | Lecythidaceae | CA | T | D | 11 |
| 13. | Carissa carandas L. | Apocynaceae | CC | S | E | 15 |
| 14. | Casearia tomentosa Roxb. | Salicaceae | BO | S | D | 12 |
| 15. | Cassine glauca (Rottb.) Kuntze | Celastraceae | CG | T | E | 14 |
| 16. | Catunaregam spinosa (Thunb.) Tirveng. | Rubiaceae | CS | T | D | 30 |
| 17. | Celtis timorensis Span. | Cannabaceae | CE | T | D | 15 |
| 18. | Colebrookea oppositifolia Sm. | Lamiaceae | CO | S | D | 18 |
| 19. | Dimorphocalyx glabellus var. lawianus (Hook.f.) Chakrab. & N.P.Balakr. | Euphorbiaceae | DL | T | E | 15 |
| 20. | Diospyros montana Roxb. | Ebenaceae | DM | T | D | 55 |
| 21. | Diospyros sylvatica Roxb. | Ebenaceae | DS | T | E | 24 |
| 22. | Diploclisia glaucescens (Blume) Diels | Menispermaceae | NA | L | E | 15 |
| 23. | Dysoxylum gotadhora (Buch.-Ham.) Mabb. | Meliaceae | DB2 | T | E | 13 |
| 24. | Elaeagnus conferta Roxb. | Elaeagnaceae | EC | L | E | 14 |
| 25. | Embelia basaal (Roem. & Schult.) A.DC. | Primulaceae | AM | S | D | 20 |
| 26. | Embelia ribes Burm.f. | Primulaceae | ER | L | E | 26 |
| 27. | Ficus nervosa B.Heyne ex Roth | Moraceae | FN | T | E | 15 |
| 28. | Ficus racemosa L. | Moraceae | FR | T | D | 15 |
| 29. | Ficus tsjahela Burm. f. | Moraceae | FT | T | D | 11 |
| 30. | Flacourtia indica (Burm.f.) Merr. | Salicaceae | FI | T | D | 25 |
| 31. | Garcinia indica (Thouars) Choisy | Clusiaceae | GI | T | E | 15 |
| 32. | Garcinia talbotii Raizada ex Santapau | Clusiaceae | PH | T | E | 27 |
| 33. | Glochidion hohenackeri (Müll.Arg.) Bedd. | Phyllanthaceae | GH | S | E | 18 |
| 34. | Gnetum ula Brongn. | Gnetaceae | GU | L | E | 28 |
| 35. | Gnidia glauca (Fresen.) Gilg | Rutaceae | LE | T | D | 18 |

Table S1 continued...

| | Species | Family | Code | Life | Leaf | n |
|---|---|---|---|---|---|---|
| 36. | Grewia tiliifolia Vahl | Malvaceae | DH | T | D | 14 |
| 37. | Heterophragma quadriloculare (Roxb.) K.Schum. | Bignoniaceae | HF | T | D | 15 |
| 38. | Heynea trijuga Roxb. ex Sims | Meliaceae | PA | T | E | 15 |
| 39. | Ixora nigricans R.Br. ex Wight & Arn. | Rubiaceae | PS | S | D | 15 |
| 40. | Jasminum malabaricum Wight | Oleaceae | JM | L | E | 15 |
| 41. | Lagerstroemia parviflora Roxb. | Lythraceae | LP | T | D | 16 |
| 42. | Leea indica (Burm. f.) Merr. | Vitaceae | LI | S | E | 11 |
| 43. | Lepisanthes tetraphylla Radlk. | Sapindaceae | LT | T | E | 15 |
| 44. | Litsea josephi S.M.Almeida | Lauraceae | LS | T | E | 21 |
| 45. | Macaranga peltata (Roxb.) Müll.Arg. | Euphorbiaceae | CH | T | E | 28 |
| 46. | Mallotus resinosus (Blanco) Merr. | Euphorbiaceae | TA | S | E | 12 |
| 47. | Mallotus philippensis (Lam.) Müll.Arg. | Euphorbiaceae | MP | T | E | 27 |
| 48. | Mangifera indica L. | Anacardiaceae | MI | T | E | 16 |
| 49. | Maytenus rothiana Lobr.-Callen | Celastraceae | BV | S | E | 19 |
| 50. | Memecylon umbellatum Burm. f. | Melastomataceae | MU | T | E | 43 |
| 51. | Meyna spinosa Roxb. ex Link | Rubiaceae | VS | T | D | 15 |
| 52. | Murraya koenigii (L.) Spreng. | Rutaceae | CP | T | D | 7 |
| 53. | Myristica dactyloides Gaertn. | Myristicaceae | MD | T | E | 11 |
| 54. | Olea dioica Roxb. | Oleaceae | OD | T | E | 14 |
| 55. | Oxyceros rugulosus (Thwaites) Tirveng. | Rubiaceae | PY | L | E | 15 |
| 56. | Pavetta indica L. | Rubiaceae | PI | S | D | 15 |
| 57. | Piper trichostachyon (Miq.) C. DC. | Piperaceae | PP | L | E | 13 |
| 58. | Pittosporum wightii A.K.Mukh. | Pittosporaceae | VI | T | E | 13 |
| 59. | Premna coriacea C.B.Clarke | Lamiaceae | PC | L | D | 15 |
| 60. | Psydrax dicoccos Gaertn. | Rubiaceae | CD | T | E | 15 |
| 61. | Rourea minor (Gaertn.) Alston | Connaraceae | RS | L | E | 15 |
| 62. | Smilax ovalifolia Roxb. ex D.Don | Smilacaceae | GT | L | D | 14 |
| 63. | Sterculia guttata Roxb. ex G.Don | Sterculiaceae | KU | T | D | 15 |
| 64. | Strobilanthes callosa Nees | Acanthaceae | KA | S | D | 13 |
| 65. | Symplocos beddomei C.B.Clarke | Symplocaceae | SB | T | E | 15 |
| 66. | Syzygium cumini (L.) Skeels | Myrtaceae | SC | T | E | 16 |
| 67. | Syzygium gardneri Thwaites | Myrtaceae | SG | T | E | 15 |
| 68. | Terminalia bellirica (Gaertn.) Roxb. | Combretaceae | TB | T | D | 8 |
| 69. | Terminalia chebula Retz. | Combretaceae | TC | T | D | 15 |
| 70. | Terminalia tomentosa Wight & Arn. | Combretaceae | SA | T | D | 15 |
| 71. | Ventilago bombaiensis Dalzell | Rhamnaceae | VB | L | E | 15 |
| 72. | Woodfordia fruticosa (L.) Kurz | Lythraceae | DA | S | D | 17 |
| 73. | Xantolis tomentosa (Roxb.) Raf. | Sapotaceae | XT | T | E | 15 |
| 74. | Zanthoxylum rhetsa DC. | Rutaceae | KH | T | D | 13 |
| 75. | Ziziphus rugosa Lam. | Rhamnaceae | TH | L | D | 14 |

**Table S2:** Relationship between measures of deciduousness and species mean flushing and senescing dates. Results presented are for $r^2$ for circular-linear correlations, and stars indication the level of significance, and * depicts $p < 0.05$, ** depicts $p < 0.01$, and *** depicts $p < 0.001$.

| Traits | Flushing date | | Senescence date | |
|---|---|---|---|---|
| Magnitude of canopy loss (MCL) | 0.7113 | *** | 0.1804 | * |
| Duration of deciduousness (DD) | 0.5270 | *** | 0.1413 | * |
| Average canopy loss (ACL) | 0.7086 | *** | 0.0891 | NS |

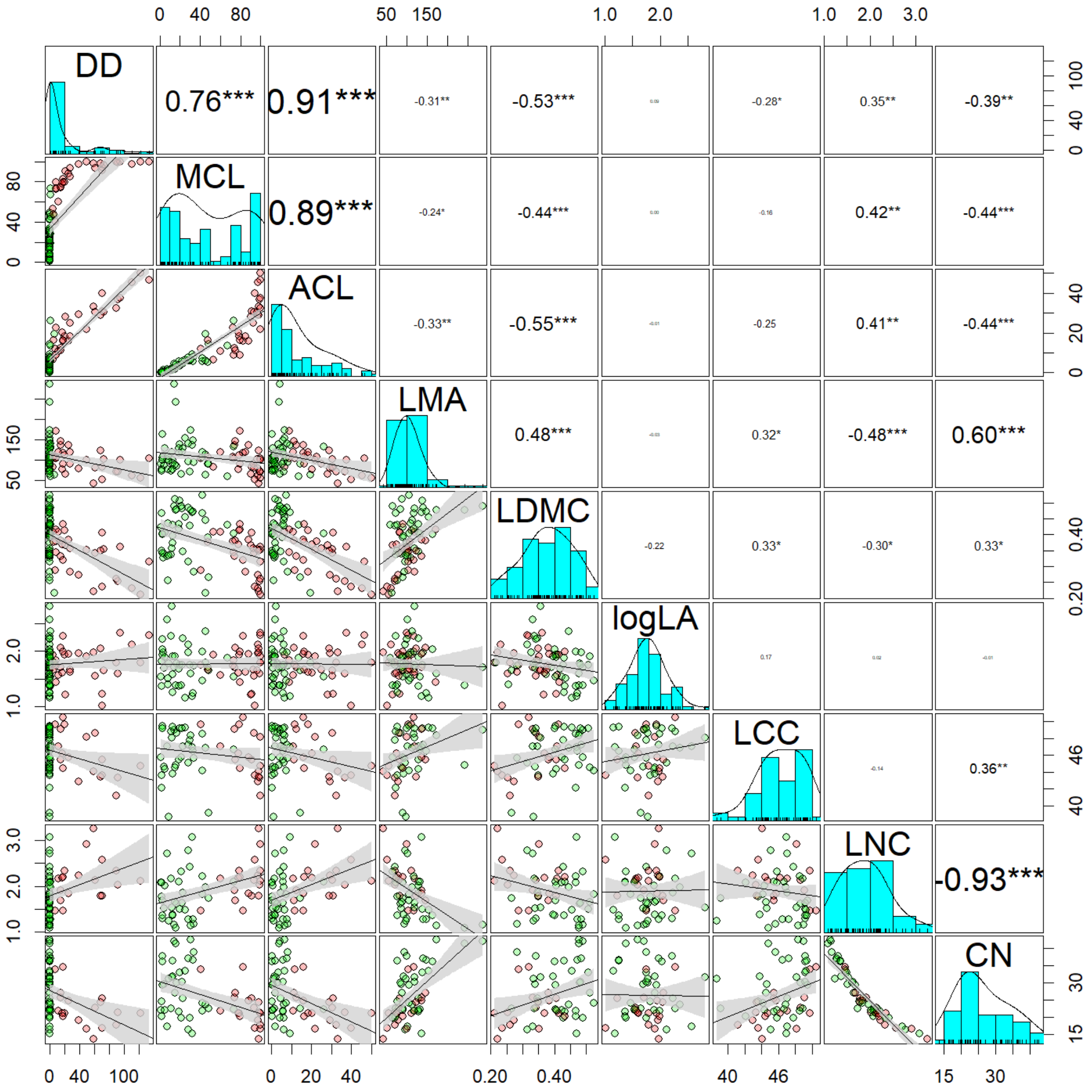


**Figure S1:** Correlation matrix for quantitative measures of deciduousness: maximum canopy loss (MCL), duration of deciduousness (DD), average canopy loss (ACL); and, leaf functional traits: leaf mass per area (LMA), leaf dry matter content (LDMC), Log transformed leaf area (logLA), leaf carbon content (LCC), leaf nitrogen content (LNC) and leaf C:N ration (CN). The pearson's correlation coefficient are shown with stars indication the level of significance, and * depicts $p < 0.05$, ** depicts $p < 0.01$, and *** depicts $p < 0.001$. Best fit lines with a 95% confidence interval band in grey are presented as a visual aid. Deciduous species are depicted in red and evergreen species in green.
